# LazyPair: scalable prediction of protein-protein interactions and interaction types

**DOI:** 10.1101/2022.02.21.481370

**Authors:** Chun Shen Lim, Bikash K. Bhandari, Paul P. Gardner

**Affiliations:** Department of Biochemistry, School of Biomedical Sciences, University of Otago, Dunedin, New Zealand; Biomolecular Interaction Centre, University of Canterbury, Christchurch, New Zealand

## Abstract

**Motivation:** Almost all cellular processes require protein-protein interactions. Common interaction types include binding, post-translational modifications, and catalysis. However, existing prediction tools do not take these interaction types into account and do not scale well on proteome-wide prediction.

**Results:** Here we show that a random forest classifier trained on per-residue physicochemical and biochemical properties is useful for predicting protein-protein interactions. Counterintuitively, we find that training random forests by individual interaction types improves accuracy. Furthermore, a combination of these specialised classifiers improves generalisability. We call our protein-protein interaction prediction tool LazyPair. More importantly, LazyPair outperforms the state-of-the-art in accuracy, generalisability and scalability.

**Availability and implementation:** LazyPair and the source code and data for reproducing our analysis are freely available at https://github.com/Gardner-BinfLab/PPI_Analysis_2022 and https://doi.org/10.5281/zenodo.6071630. The web server version and the source code are freely available at https://tisigner.com/lazypair/ and https://github.com/Gardner-BinfLab/TISIGNER-ReactJS, respectively.

## INTRODUCTION

Protein-protein interactions (PPIs) are involved in nearly all cellular processes and metabolisms. Common experimental methods for detecting PPIs include yeast two-hybrid assays, anti-tag coimmunoprecipitation, and tandem affinity purification. These experimentally validated PPIs are curated and maintained by a number of databases, e.g., BIOGRID (Oughtred *et al*., 2021), BioPlex (Huttlin *et al*., 2021), DIP (Salwinski *et al*., 2004), IntAct (Orchard *et al*., 2014), mentha (Calderone *et al*., 2013), MINT (Licata *et al*., 2012), signor (Licata *et al*., 2020), and STRING (Szklarczyk *et al*., 2021). However, these curated PPIs are mostly PPIs detected in human and yeast cells; there is a need to predict PPIs in other species.

Prediction of PPIs is a non-trivial task. Recent tools (Chen *et al*., 2019; Hashemifar *et al*., 2018) have been found to lack generalisability in predicting PPIs across species (Sledzieski *et al*., 2021). Furthermore, most tools were trained and evaluated using specific studies/databases, raising a question of whether these tools are generalisable across studies/databases. For instance, a tool predominantly trained on PPIs found in yeast two-hybrid assays may be suboptimal in predicting PPIs found in affinity purification-based assays. This is because weak and transient interactions are more likely to be detected in yeast two-hybrid assays than affinity purification-based assays. In addition, existing tools are not scalable for proteome-wide prediction. With these in mind, we aim to develop an accurate, generalisable and scalable PPI prediction tool.

## METHODS

### Data

We retrieved PPIs from D-SCRIPT (Sledzieski *et al*., 2021) and eight databases, namely BIOGRID 4.4.204 (Oughtred *et al*., 2021), BioPlex 3.0 (Huttlin *et al*., 2021), DIP 20170205 (Salwinski *et al*., 2004), IntAct 2021-10-13 (Orchard *et al*., 2014), mentha 2021-12-20 (Calderone *et al*., 2013), MINT 2021-12-19 (Licata *et al*., 2012), signor Jan2022 release (Licata *et al*., 2020), and STRING 11.5 (Szklarczyk *et al*., 2021). We selected physical-only PPIs from BIOGRID and STRING.

For specific interaction types, we used the annotations in signor and STRING 10.5 (https://version-10-5.string-db.org/download/protein.actions.v10.5/9606.protein.actions.v10.5.txt.gz).

### Machine learning

We calculated a total of 553 features from protein sequences using the aaindex1 function of protlearn v0.0.3 (Tadorfer, 2021; Kawashima *et al*., 2008). AAindex1 is a set of per-residue values that capture different physicochemical and biochemical properties of 20 standard amino acids. Protlearn is a tool to process, calculate and select features from protein sequences. For each protein pair, we took the average value for each feature. We then used these features to train and evaluate random forests using scikit-learn v1.0.2 (Pedregosa *et al*., 2011).

### Benchmarking

To assess the generalisability of each method, 1000 positive and 1000 negative PPIs were sampled 20 times from each PPI database and low probability PPIs (pInt <0.1 in BioPlex 3.0), respectively. The redundant PPIs were removed—for some pairs (33.5%) D-SCRIPT did not produce a result, in order to generate a fair benchmark these were not included in the comparison with LazyPair. AUROC (area under the receiver operating characteristic curve) and AUPRC (area under the precision-recall curve) were computed from 100 bootstrap resamplings of the predicted results, where each sample contained 500 positives and 5000 negatives. In reality, the number of protein pairs that do not interact are likely to be much more frequent than pairs that do, hence our imbalanced positive and negative sets.

To evaluate scalability, PPI prediction tools were run three times using a single process without GPUs on a high performance computer (Red Hat Enterprise Linux 7.9, 2x 64-core AMD EPYC 7702, 1023GiB memory), and timed using /usr/bin/time-f ‘%E’ <command>.

### Statistical analysis

Data analysis was done using Pandas v1.2.4 (McKinney, 2010), numpy v1.20.1 (van der Walt *et al*., 2011), scipy v1.6.2 (Virtanen *et al*., 2020), and scikit-learn v1.0.2 (Pedregosa *et al*., 2011). Plots were generated using Matplotlib v3.3.4 (Hunter, 2007), Seaborn v0.11.1 (Waskom *et al*., 2014), and UpSetPlot v0.6.0 (Nothman, 2021).

### Code and data availability

LazyPair and Jupyter notebooks of our analysis can be found at https://github.com/Gardner-BinfLab/PPI_Analysis_2022 and https://doi.org/10.5281/zenodo.6071630. The source code for the LazyPair web application can be found at https://github.com/Gardner-BinfLab/TISIGNER-ReactJS (Bhandari *et al*., 2021).

## RESULTS AND DISCUSSION

### Physicochemical and biochemical properties of amino acids are useful for predicting PPIs

We sought to develop an accurate and generalisable PPI prediction tool for proteome-wide analysis. Therefore, we extracted features from protein sequences using AAindex1 that represents the physicochemical and biochemical properties of amino acids (Kawashima *et al*., 2008), and trained a random forest for PPI prediction (called Lazy_0_). We used the same datasets that were recently used to develop the state-of-the-art prediction tool D-SCRIPT (Sledzieski *et al*., 2021). In particular, we used human PPIs for training and PPIs in other model species for evaluation, including *Mus musculus, Drosophila melanogaster, Caenorhabditis elegans, Saccharomyces cerevisiae, Escherichia coli*. These positive sets were physical-only PPIs originally from STRING 11 (Szklarczyk *et al*., 2021). The negative sets were protein pairs previously generated by a random sampling approach (Sledzieski *et al*., 2021; Hashemifar *et al*., 2018).

The above steps allow us to directly compare the performance results of Lazy_0_ with previously published results for D-SCRIPT (Sledzieski *et al*., 2021) and PIPR (Chen *et al*., 2019). In general, Lazy_0_ outperformed PIPR but underperformed D-SCRIPT (Table 1).

**Table 1.**
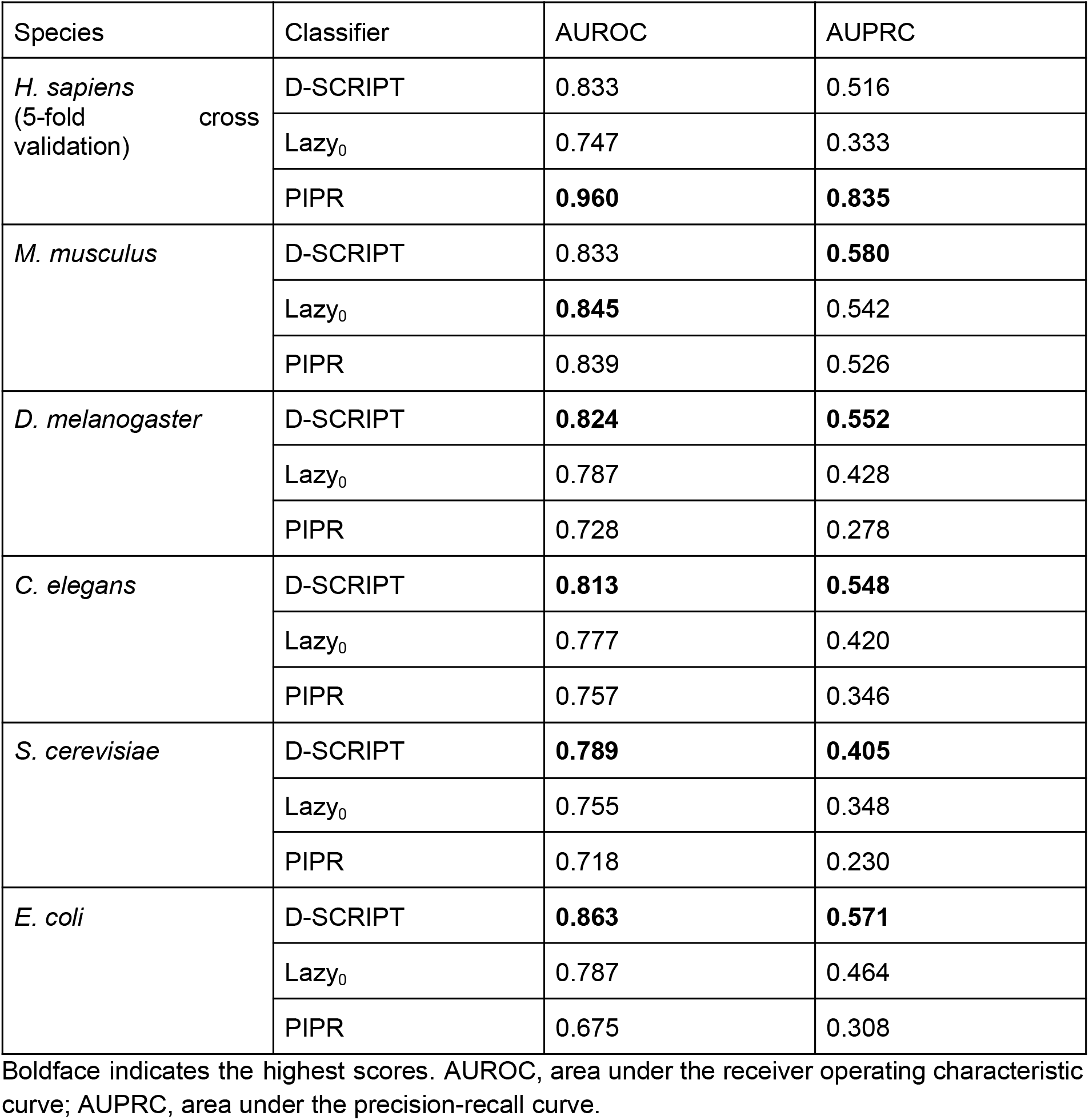
Performance results for classifiers trained on protein-protein interactions (PPIs) in humans. The performance results for PIPR (Chen *et al*., 2019) and D-SCRIPT were taken from a previous study (Sledzieski *et al*., 2021). Both the training and test data sets for the below model organisms were derived from a previous study (Sledzieski *et al*., 2021).

Next, we asked how generalisable the prediction tools are across PPIs in different databases. Interestingly, Lazy_0_ outperformed D-SCRIPT in five out of eight instances (Fig 1). However, the performance results presented in Fig 1 that are for PPIs from different databases were approximately 20% lower for both methods than for the model organism results presented in Table 1. This is likely due to overfitting on human PPIs taken from the STRING database. A closer look revealed that most PPIs were unique across databases (Fig 2). Furthermore, an alternative negative test set was used for Fig 1, i.e., low probability PPIs in BioPlex 3.0 (Huttlin *et al*., 2021), rather than randomly sampled protein pairs as above.

**Fig 1.**
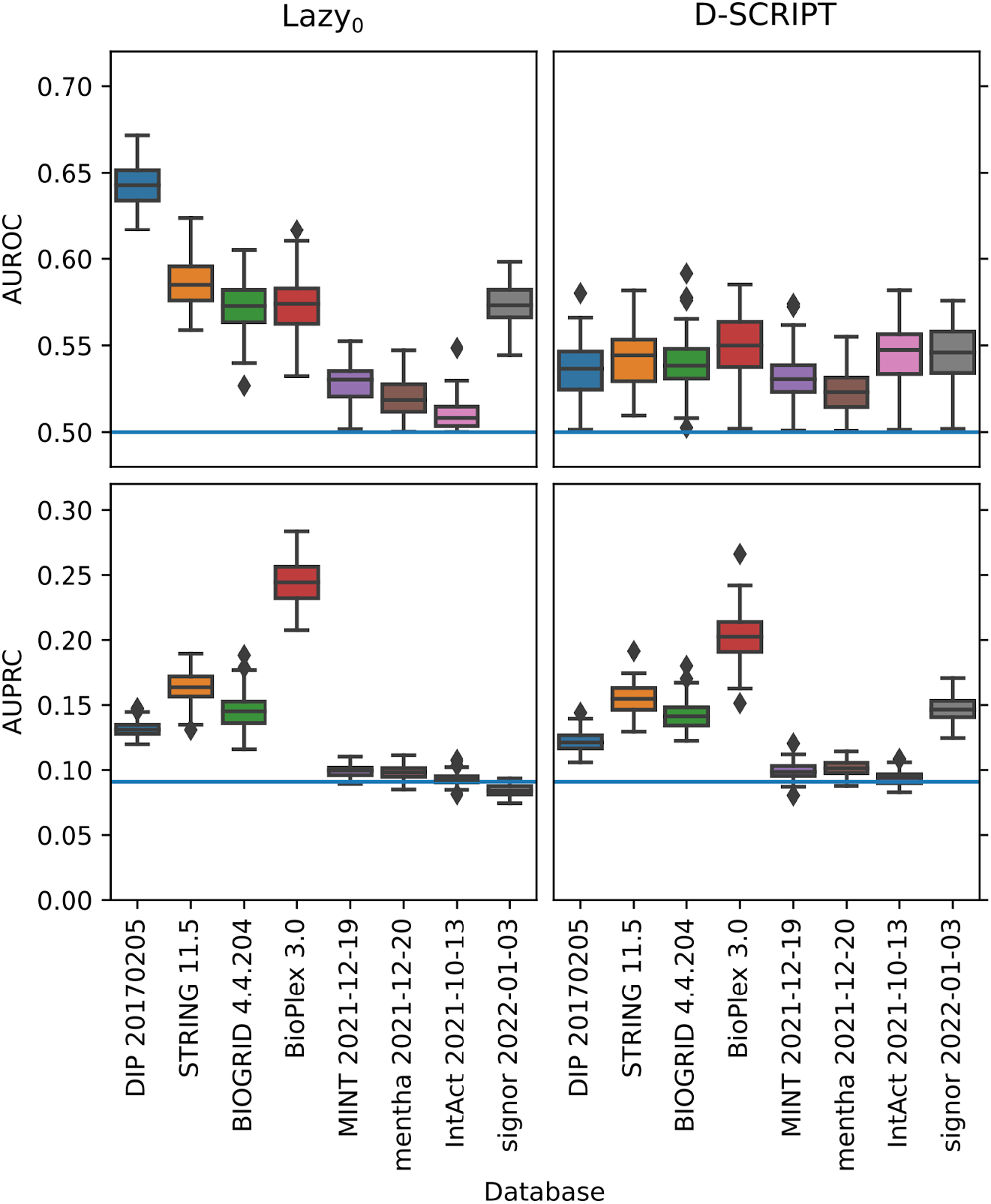
Random forest (Lazy_0_) trained on physical protein-protein interactions (PPIs) in humans is more generalisable across PPI databases than that of convolutional neural networks. Low probability PPIs were used as the negative test set (pInt <0.1 in BioPlex 3.0). The performance results of Lazy_0_ and D-SCRIPT were computed from 100 bootstrap replicates. The blue lines indicate the performance results of random classifiers. AUPRC, area under the precision-recall curve; AUROC, area under the receiver operating characteristic curve.

**Fig 2.**
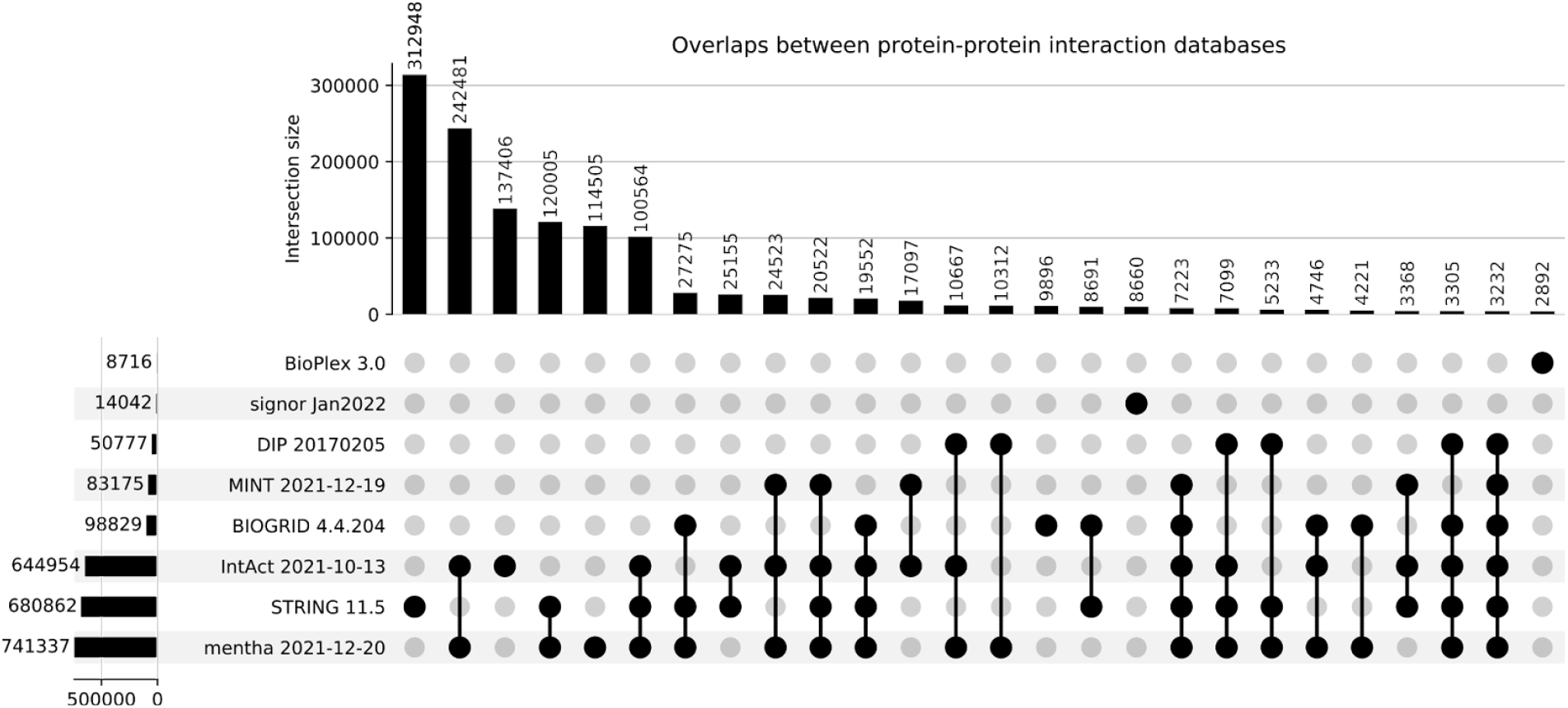
Most PPIs are unique across databases. Physical-only PPI releases were used whenever possible, as for BIOGRID and STRING.

**Fig 3.**
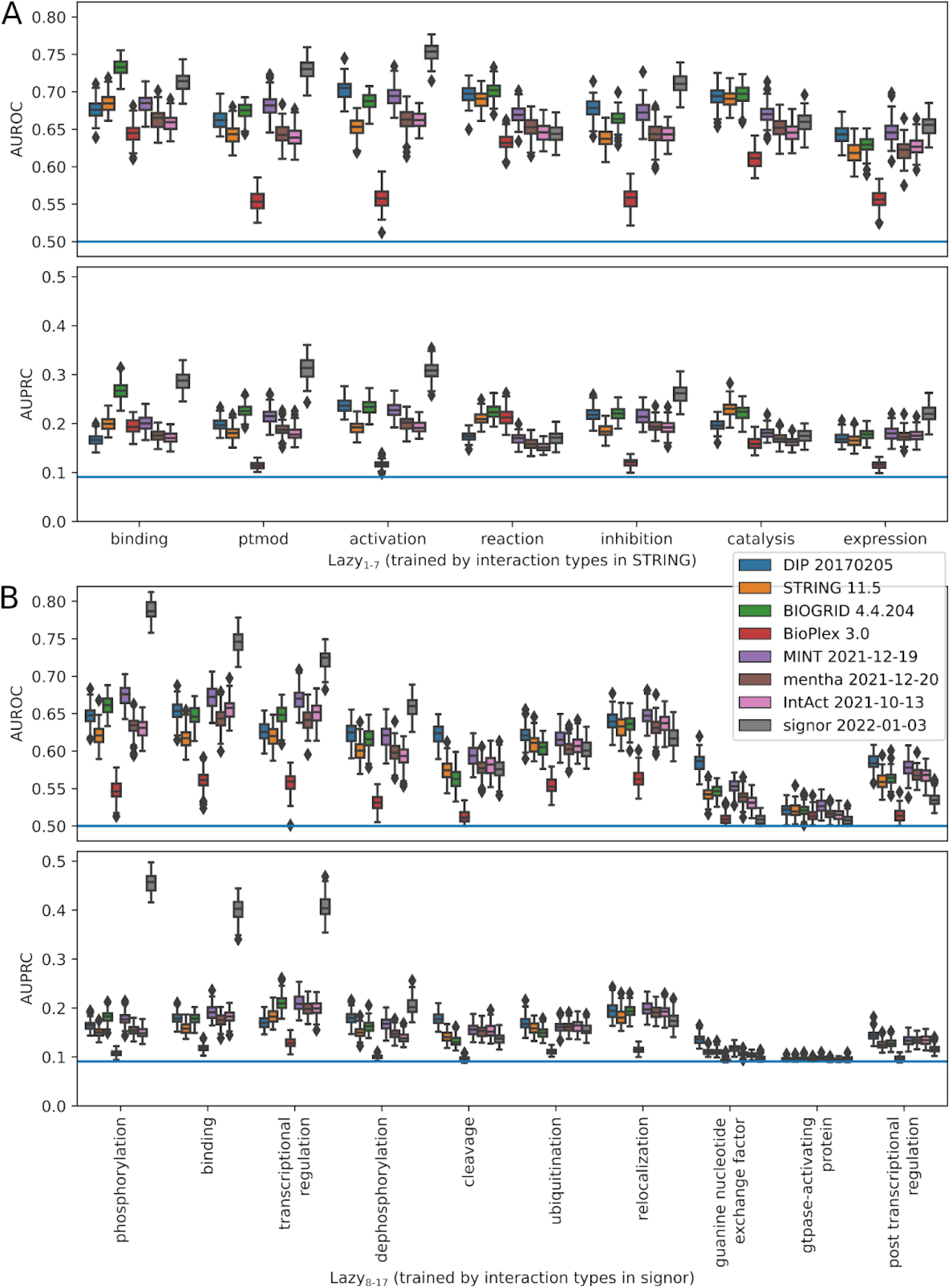
Random forests (Lazy_1-17_) trained by specific interaction types show improved performance results. **(A)** All 7 interaction types in STRING 10.5 were used for training, including binding (n=15,925), ptmod (post-translational modification, n=926), activation (n=1,283), reaction (n=7,295), inhibition (n=542), catalysis (n=3,525), expression (n=265). **(B)** The 10 largest interaction types in signor were used for training, including phosphorylation (n=2,554), binding (n=2,153), transcriptional regulation (n=2,484), dephosphorylation (n=254), cleavage (n=156), ubiquitination (n=65), relocalization (n=138), guanine nucleotide exchange factor (n=27), gtpase-activating protein (n=29), post transcriptional regulation (n=51). For negative PPIs, two independent sets of proteins with low probability PPIs were used for training and evaluation (pInt <0.1 in BioPlex 3.0). The performance results of Lazy_1-17_ were computed from 100 bootstrap resampling. The blue lines indicate the performance results of random classifiers. AUPRC, area under the precision-recall curve; AUROC, area under the receiver operating characteristic curve.

### Random forests trained by specific interaction types show improved accuracy and generalisability

We reasoned that the generalisability issue may be amplified by the complexity of protein-protein interaction types. As some interaction types are more transient than others, e.g. phosphorylation versus binding/assembly, transient and stable PPIs may be attributed to distinct sequence properties. Therefore, we divided the PPIs in STRING and signor by interaction types and separately trained random forests on these subsets. We did not use the interaction types in other databases as only high-level interaction types were available, i.e., association (mi:0914), physical association (MI:0915), and direct interaction (MI:0407).

Interestingly, most of the specialised classifiers outperformed the generic classifier (Fig 3 versus Fig 1, Lazy_1-10_ versus Lazy_0_). We ranked the features by Gini importance and found that a “hydrophobicity coefficient” (WILM950103) is among the strongest features for these classifiers. There is a strong correlation between feature strengths for “binding”, “catalysis” and “reaction” interaction types—the sequence features most strongly associated with these are hydrophobicity, alpha and turn propensities, and an “other” class of AAindex1 (Supplementary Figs S1). However, many of these specialised classifiers also have distinct feature rankings, suggesting that proteins involved in different interaction types are distinct (Supplementary Figs S1 and S2, Spearman’s correlation coefficients ~0.5).

We also found that the median probability of the specialised classifiers improved generalisability (Fig 4). We then repeated the training step using the complete datasets/subsets and combined these classifiers as a PPI prediction tool called LazyPair.

**Fig 4.**
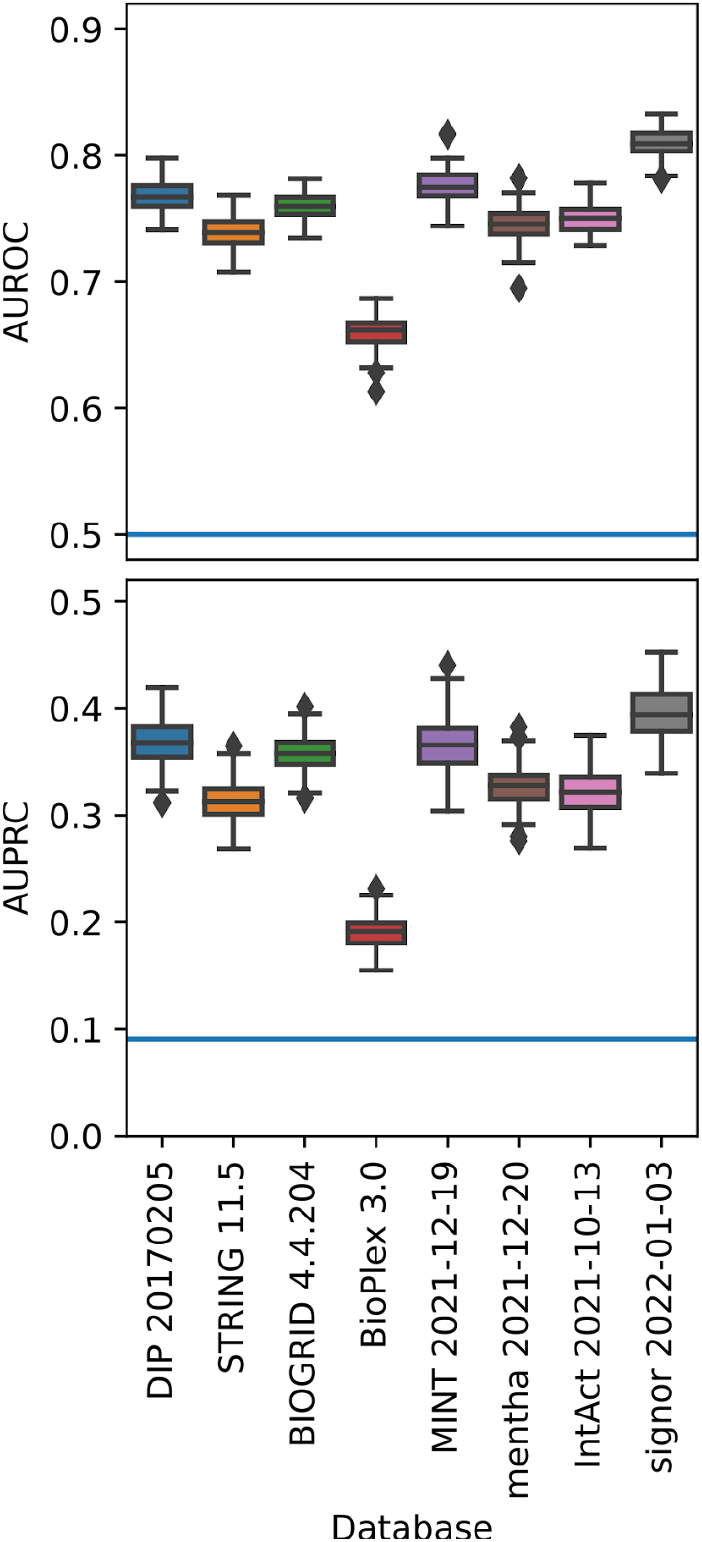
median probabilities of Lazy_0-17_ improve generalisability. The performance results were computed from 100 bootstrap resampling. The blue lines indicate the performance results of random classifiers. AUPRC, area under the precision-recall curve; AUROC, area under the receiver operating characteristic curve.

**Fig 4.**
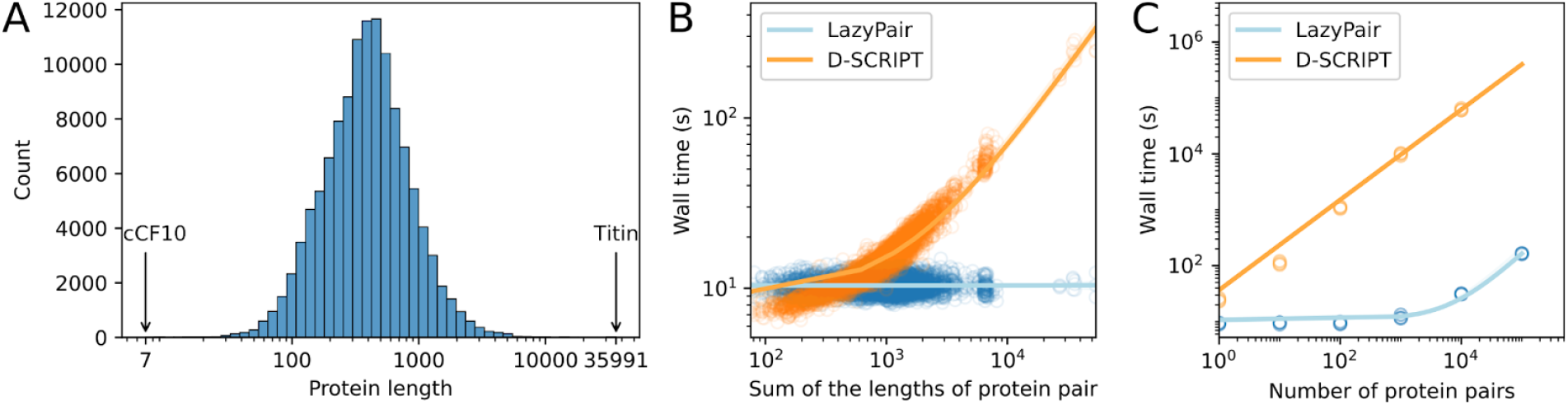
LazyPair is scalable for large-scale PPI prediction. (A) Distribution of the lengths of protein sequences from PPI databases. The shortest sequence is the 7-residue bacterial sex pheromone cCF10, whereas the longest sequence is the 35,991-residue human titin. (B) Wall time of PPI prediction tools for single protein pairs. LazyPair took approximately *O*(*n*) time with increase in length of the interacting pairs. (C) Wall time of PPI prediction tools for different sample sizes.

### LazyPair is scalable for proteome-wide PPI prediction

For scalability benchmarking, we tested the effects of (i) protein sequence length and (ii) sample size. LazyPair outperformed the state-of-the-art prediction tool in both tests (Fig 4).

In the first test, we first selected single sequences spanning across the distribution of sequence lengths from PPI databases (Fig 4A). We then retrieved the interacting partners and tested the PPI prediction tools three times (Fig 4B). LazyPair took approximately *O*(*n*) with negligible wall time as the features were simply the arithmetic means of per-residue values (AAindex1).

In the second test, we performed random sampling by increasing the sample size of PPI test sets. This allowed us to obtain the test sets in ascending order of magnitude and further test the PPI prediction tools three times (Fig 4B). LazyPair only showed signs of slowing down at 1,000 protein pairs (12.2 ± 1.0 s versus 2 h 44 min 10.7 ± 9 min 3.4 s, using a single process without GPUs; 2×64-core AMD EPYC 7702, 1023GiB memory).

To attain proteome-wide PPI prediction, we have implemented multiprocessing in LazyPair. We found that LazyPair took 2 h 17 min 39.7 s ± 8 min 21.4s to predict all 9,644,832 possible PPIs in *E. coli* (UniProt: UP000000625; 4392 proteins) using 128 parallel processes.

In conclusion, we have developed an accurate and generalisable PPI prediction tool that is capable of predicting proteome-wide PPIs. To make LazyPair accessible to a broader audience, we have also developed a web version available at https://tisigner.com/lazypair.

## ACKNOWLEDGEMENTS

This work was supported by the Royal Society of New Zealand Te Apārangi [Marsden grant: 19-UOO-040 to P.P.G.] and the Ministry of Business, Innovation and Employment [MBIE Data Science Programmes grant: UOAX1932 to P.P.G.].

## AUTHOR CONTRIBUTIONS

C.S.L. and P.P.G. conceived the study; C.S.L. analysed the data and developed LazyPair; B.K.B. developed the LazyPair web application; C.S.L. drafted the manuscript; C.S.L., B.K.B., and P.P.G. reviewed, edited, and approved the manuscript.

