## Supplementary material for "LazyPair: scalable prediction of protein-protein interactions and interaction types": Fig S1, Fig S2

### SUPPLEMENTARY MATERIALS

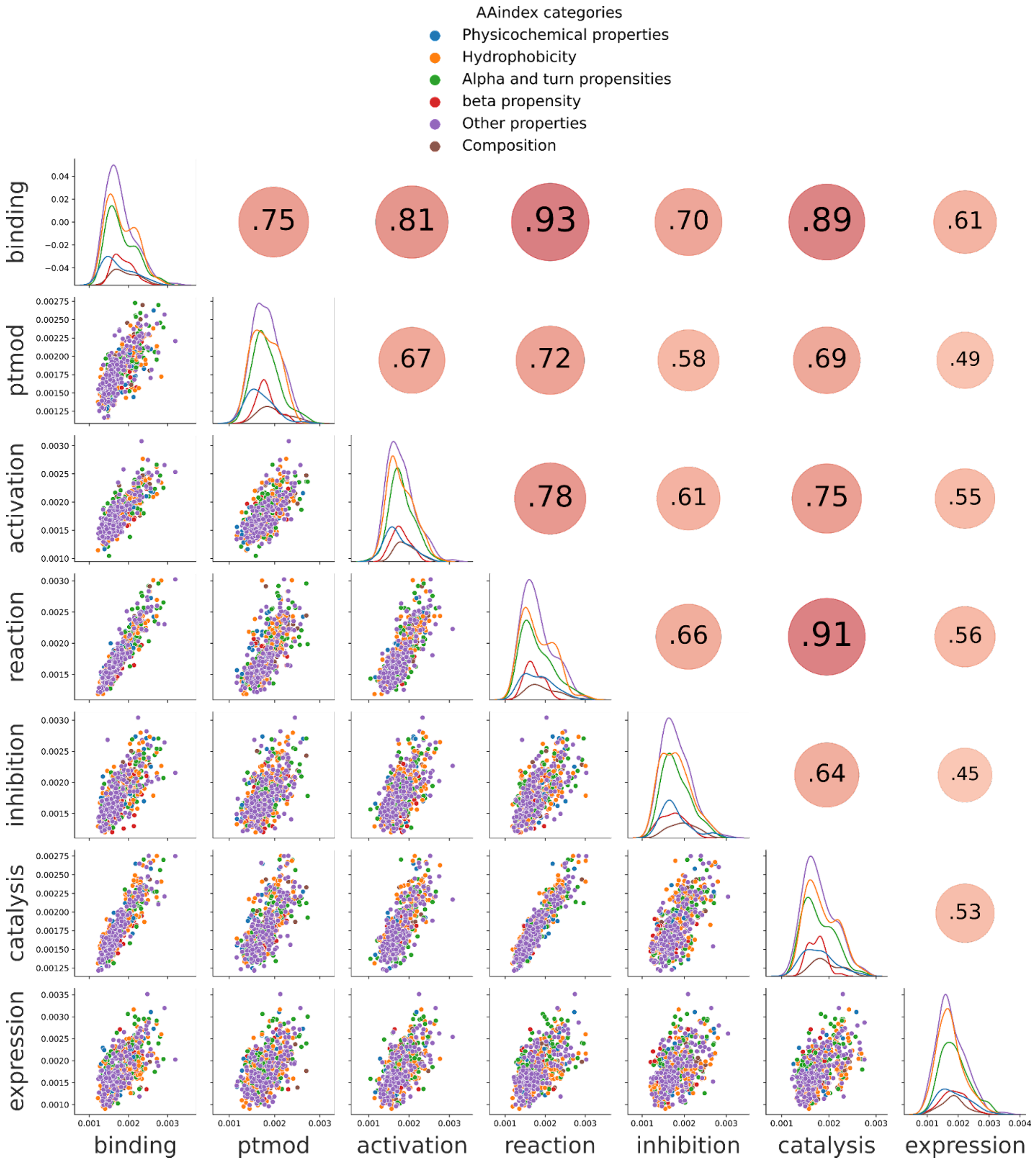


**Fig S1. Correlation matrix plot of the feature importances of the final random forest classifiers trained on 7 STRING interaction types (Lazy$_{1\text{-}7}$).** A total of 553 features were extracted using AAindex1 ('Other properties' include 151 new features that have not been categorised). These STRING interaction types include binding (n=15,925), ptmod (post-translational modifications, n=926), activation (n=1,283), reaction (n=7,295), inhibition (n=542), catalysis (n=3,525), expression (n=265). Low probability PPIs were used as negatives (pInt <0.1 in BioPlex 3.0, n=1,323,898).

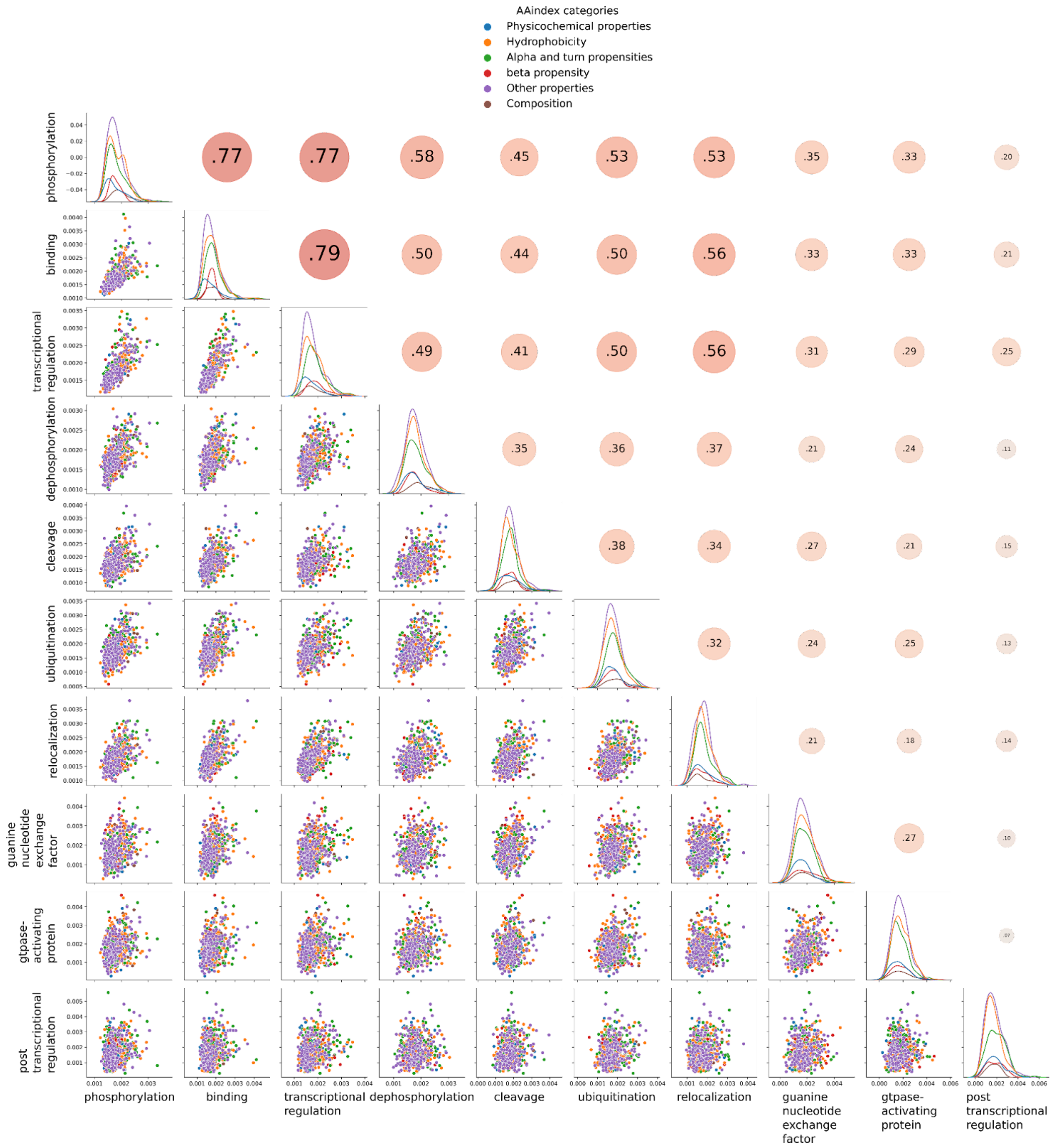


**Fig S2. Correlation matrix plot of the feature importances of the final random forest classifiers trained on the 10 largest signor interaction types (Lazy$_{8\text{-}17}$).** A total of 553 features were extracted using AAindex1 (‘Other properties’ include 151 new features that have not been categorised). These signor interaction types include phosphorylation (n=4,542), binding (n=4,379), transcriptional regulation (n=2,736), dephosphorylation (n=520), cleavage (n=234), ubiquitination (n=272), relocalization (n=290), guanine nucleotide exchange factor (n=93), gtpase-activating protein (n=87), post transcriptional regulation (n=57). Low probability PPIs were used as negatives (pInt <0.1 in BioPlex 3.0, n=1,323,898).
